# Multiplexed enrichment and genomic profiling of peripheral immune cell subsets on a microfluidic chip

**DOI:** 10.1101/261321

**Authors:** Miguel Reyes, Dwayne Vickers, Kianna Billman, Thomas Eisenhaure, Paul Hoover, Edward Browne, Deepak A. Rao, Nir Hacohen, Paul C. Blainey

**Affiliations:** Broad Institute of MIT and Harvard, Cambridge, MA, USA.; Department of Biological Engineering, Massachusetts Institute of Technology, Cambridge, MA, USA.; Division of Rheumatology, Immunology, Allergy, Brigham and Women’s Hospital and Harvard Medical School, Boston, MA, USA.; Center for Cancer Research, Department of Medicine, Massachusetts General Hospital, Boston, MA, USA

## Abstract

The human immune system consists of many specialized cell subsets that simultaneously carry out a diverse range of functions using overlapping pathways and signals. Subset-specific immune profiling can resolve immune activity in autoimmune disease, cancer immunity, and infectious disease that may not be discoverable or detectable in analyses of crude blood samples. The activity of specific subsets may help predict the course of disease and response to therapy in certain patient populations. Here, we present a low-input microfluidic system for sorting immune cells into subsets and profiling their cellular states by gene expression analysis using full-length RNA-seq. Our system is robust and has the potential to make multiplexed subset-specific analysis routine in many research laboratories and clinical settings. We validate the device’s technical performance by benchmarking its subset enrichment and genomic profiling performance against standard protocols. We make the added value of subset-resolved profiling over crude samples clear through *ex vivo* experiments that show subset-specific stimulated responses. Finally, we demonstrate the scalability of our device by profiling four immune cell subsets in blood from systemic lupus erythematosus (SLE) patients and matched controls enrolled in a clinical study. The results from our initial cohort confirm the role of type I interferons in lupus pathogenesis and further show that the canonical interferon signature for SLE is prominent in B cells, demonstrating the ability of our integrated analytical platform to identify cell-specific disease signatures.

## INTRODUCTION

Millions of immune cells can be obtained from a small blood draw, yet most clinical methods for immune profiling fail to resolve the biological information contained within these cells. Recently, profiling the immune state of individuals using gene expression analysis of peripheral blood mononuclear cells (PBMCs) has become instrumental in defining immune signatures and disease states in humans. These studies provide insight into the mechanisms of complex immune responses that occur in infection^1,2^ and autoimmunity^3–5^, which are difficult to recapitulate in murine models^6–8^. Furthermore, expression signatures can be used to stratify individuals into different disease subtypes^9–13^ or predict individualized clinical prognoses^14–16^. More recently, gene expression profiles from specific cell types or cell “subsets” were shown to be better discriminants of immune status than bulk PBMC profiles due to the diversity of leukocyte responses^17–19^. In addition, new immune subsets and cellular states, some of which are indicative of impaired immune function, have been discovered through gene expression profiling of PBMCs at the single cell level^20–23^. Such observations have stirred interest in probing the expression and monitoring the activity of these subsets in particular. As a whole, this body of work suggests that molecular profiling of PBMC subsets is poised to become an important tool in basic studies of immune disease as well as a clinical tool useful for predicting and monitoring patient outcomes.

Despite its potential as a tool for immunomonitoring, current methods for subset-specific expression profiling are ill-suited for large studies and clinical translation. First, technologies for cell subset enrichment such as fluorescence-activated cell sorting (FACS) are both capital intensive and require substantial attention from highly trained staff. As a result, FACS is challenging to scale for large-number samples generated from clinical studies (especially those that generate large sample sets by analyzing multiple cell subsets across patients at different time points). In addition, FACS requires a minimum sample input to establish gates for each target subset, which can constrain its application to low-quantity samples and projects targeting many subsets from each sample. Second, the throughput of RNA-seq library construction is limited by reagent cost and labor. Implementing library construction at high throughput using conventional pipetting robots is feasible, but capital intensive, inflexible, and only tractable for the largest studies and the largest clinical centers. Conventional magnetic-affinity cell sorting (MACS) has the potential to be automated, but available commercial systems are low-throughput and custom liquid handling systems suffer the same drawbacks just described for their application in RNA-seq library construction. Because of these limitations, most clinical gene expression studies are currently limited to whole-blood or total PBMC profiling^9–11,14,15^, which fails to resolve expression signatures from most cell subsets due to confounding signals from more abundant cell populations. To efficiently identify and monitor important disease signatures in lower-abundance subsets, we developed a microfluidic system that integrates both human PBMC subset enrichment and library construction for genome-wide expression measurements by RNA-seq.

The microfluidic system carries out multiplexed enrichment of target cell subsets based on affinity for cell surface markers by MACS and high-sensitivity sequence library construction for full-length RNA-seq based on Smart-seq chemistry. Using 50,000 cells as input, the device can purify multiple PBMC cell subsets with high purity while detecting up to 10,000 genes in each subset. In testing immune stimulation and challenge *ex vivo*, we highlight the importance of subset-specific profiling by showing differential responses across four selected subsets. Finally, we apply the microfluidic device to profile PBMCs of SLE patients and identified clear differences in the transcriptomic states of healthy individuals and SLE patients in multiple immune cell subsets. By integrating multiplexed enrichment and library construction workflows in a single device, our platform enables scalable PBMC sample preparation for large clinical studies and allows for both high-resolution and high-throughput profiling of the immune system. We foresee the application of this system as a routine tool in monitoring immune responses in clinical studies and a potential diagnostic tool for patients with complex and/or pressing immune conditions.

## RESULTS

### Microfluidic device design

We designed a two-layer microfluidic device capable of semi-automated cell isolation, cell disruption, and sequence library construction protocols. This system integrates microfluidic liquid handling with magnetic affinity purification. The device is fabricated using established methods for two-layer soft-lithography^24^ and contains 39 micromechanical valves controlled by an external pneumatic valve controller^25^. The device consists of three main chambers partitioned by microvalves (**Fig. 1a**), each having different capacities (1 μl, 2 μl, 4 μl). The largest chamber is rectangular in shape and is utilized for cell isolation (**Supplementary Note**), while the two smaller “rotary reactors” are used for library construction^26^. These reactors have internal microvalves that are used to formulate samples and reagents, and to mix these by peristalsis around the circular channel (**Fig. 1b**). Bead resuspension is achieved by peristalsis in the smaller rotary reactors and by a moving magnetic field in the large rectangular chamber. A 675 micron thick silicon wafer was used as the substrate for these microfluidic devices to allow rapid heat transfer during temperature changes called for in the protocol, particularly for PCR (due to its thinness and high thermal conductivity). The substrate thinness also enables small external permanent magnets to be placed in close proximity to magnetic beads in the device chambers and to subject these to strong magnetic forces. The magnets are used to move beads between chambers and hold beads in place during buffer exchange steps. With such device functionality, we are able to automate many steps in the complex protocol for cell sorting, cell disruption, and RNA-seq library construction in a simple microarchitecture consisting of just three microfluidic chambers (**Supplementary Fig. 1**). The three-chamber design element is modular and constitutes a scalable microarchitecture for devices with variable sample multiplexing capacity. The data presented here were produced using 10-channel devices, although we have fabricated devices with 6 – 30 channels. Like the devices employed in our previously published system for microbial genomic DNA sample preparation^25^, the devices described here can be reused following a simple washing procedure (particular devices were used up to 4 times in this study).

**Figure 1.**
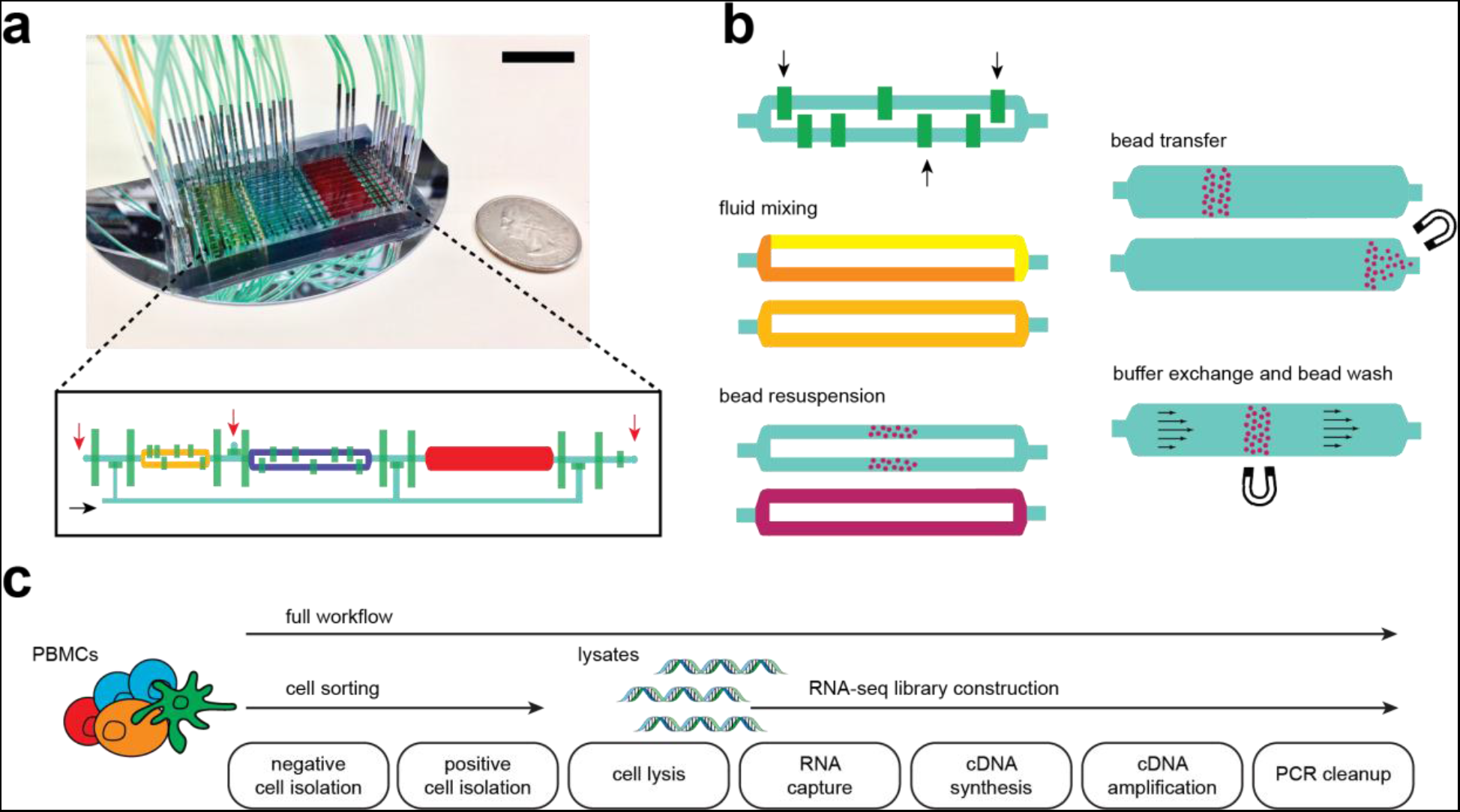
Microfluidic chip design and integrated workflows. (a) Photograph of a 10-channel chip filled with yellow, blue, and red dye to highlight compartments, control valves are filled with green dye (black bar indicates 2 cm). Inset shows diagram of one channel, where compartments and control lines are colored similar to the photograph; red and black arrows indicate sample input/output and reagent input ports, respectively. (b) Schematic showing key device capabilities that enable various sample preparation steps. Black arrows indicate mixing valves that alternately open and close to generate flow within a compartment, allowing for reagent mixing and bead resuspension without external fluid input. Permanent magnets are utilized for moving magnetic beads across different compartments or preventing their flow during buffer exchange and washes. (c) Full and partial (cell sorting and RNA-seq) sample preparation workflows implemented in the microfluidic chip.

### Microfluidic cell sorting and low-input RNA-seq

To validate the performance of our microfluidic device, we independently benchmarked the subset enrichment and RNA-seq workflows against standard protocols (**Fig. 1c**). We tested our workflows using adult peripheral PBMC samples from healthy subjects obtained from a commercial supplier (Research Blood Components) at an input level of 50,000 cells per enrichment. We first implemented MACS on the microfluidic device and configured an eight-color flow cytometry analysis as a readout of the purity and yield of the purified cell subsets (**Supplementary Fig. 2**). We optimized the conditions for microfluidic cell subset isolation by testing different reagents, incubation, and washing procedures and compared the results of the optimized protocol to conventional benchtop MACS (**Supplementary Fig. 3, Methods**). We found that the purity of the subsets isolated using the optimized microfluidic MACS protocol were about the same as those as conventional benchtop MACS, suggesting that the MACS reagents were performing up to their potential in both formats.

We tested positive selection of target cells, negative depletion of non-target cells, and sequential isolation using both modalities in tandem. Total T cells were isolated by depleting cells expressing markers for lineages other than T cells. The total T cell population was then positively selected for either CD4 or CD8 to isolate helper and cytotoxic T cell subsets separately. The prior negative selection reduced contamination from non-target lineages that express CD4 or CD8. B cells and monocytes, on the other hand, were effectively isolated using single positive selection for CD19 and CD14, respectively. The device consistently achieved good purity (80 ± 8%) and excellent yield (76 ± 21%) for multiple targets and modes of isolation (**Fig. 2a-b**), leading to 2- to 13-fold enrichment of the selected cell types, which is close to the maximum theoretical enrichment possible for the relatively abundant cell subsets tested. This result shows that microfluidic cell sorting with magnetic beads presents a viable alternative to conventional sorting approaches and demonstrates the possibility of subset-specific enrichment with limited quantity samples.

**Figure 2.**
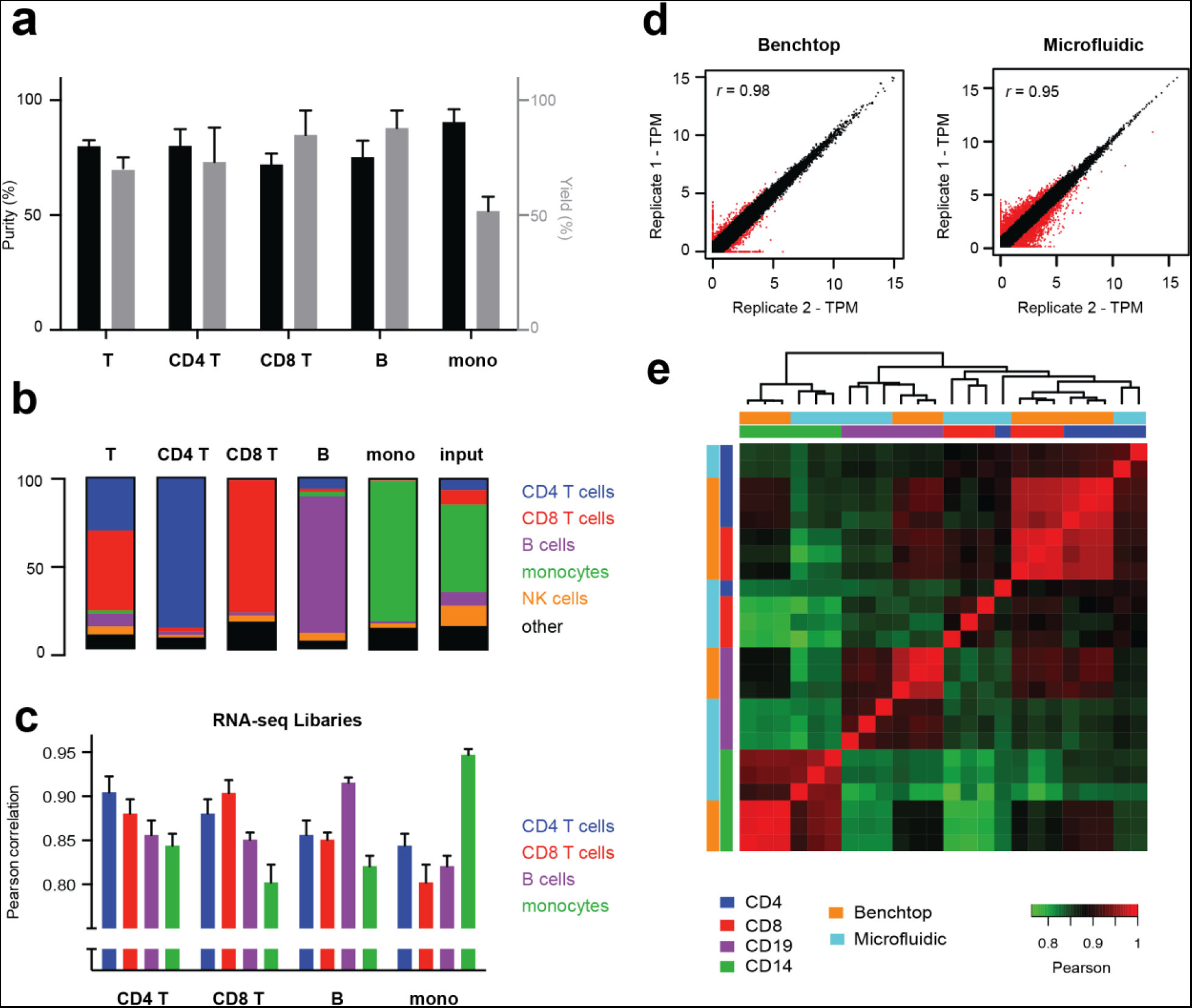
Chip performance and validation. (a-b) Representative purity (black bars), yield (gray bars), and composition of immune cell types after microfluidic sorting. Yield is determined relative to fraction of the target subset in the input sample. Error bars indicated S.E.M., n=3. (c) Pearson correlations between RNA-seq libraries of the four cell subsets processed through the microfluidic chip. (d) Scatterplot showing technical replicability of standard and microfluidic RNA-seq. Red points indicate genes with greater than 2-fold change between replicates. (e) Correlation matrix between standard and microfluidic RNA-seq libraries for four FACS-sorted cell lysates with single positive markers (CD4, CD8, CD14, and CD19).

Based on the cell isolation testing, we expected to capture thousands of cells in each subset using our microfluidic device. With these relatively low numbers in mind, we implemented a sensitive RNA-seq protocol^27^ (Smart-seq2) in the chip with minor modifications. Instead of SPRI-based clean-up for RNA extraction, we utilized custom-prepared poly-dT capture beads (**Methods**) that captured mRNA molecules in lysate by direct hybridization to enable purification and subsequent solid-phase reverse transcription. Our protocol calls for amplifying cDNA by PCR, purifying the products with SPRI, and subsequently recovering the samples from the device for Nextera fragmentation and enrichment PCR closely following the standard Smart-seq2 method. The cDNA amplicons from the microfluidic device showed the expected size distribution and the RNA-seq datasets resulting from such samples show high technical reproducibility (Pearson correlation of 0.88 ± 0.04) and correlate well with libraries produced using the standard Smart-seq2 protocol on the benchtop across for four different cell subsets (0.90 ± 0.03) (**Fig. 2d, Supplementary Fig. 4, Table 1**). Despite the similarity between the gene expression profiles of the four subsets (**Fig. 2c**), the sequence libraries produced in our workflow can distinguish the subsets based on simple correlation and clustering procedures (**Fig. 2e**). In addition, the enrichment of polyadenylated RNA in the microfluidic protocol reduced the number of ribosomal RNA reads and improved transcript mapping rates over the standard Smart-seq2 protocol (**Table 1, Supplementary Fig. 5)**. Combining RNA-seq with cell isolation in an integrated workflow yields libraries of similar quality (**Table 1, Supplementary Fig. 6**). These results demonstrate that full-length cDNA synthesis and amplification by PCR can be implemented in a microfluidic device with input from on-device-enriched cell subsets to support RNA-seq and that reduction in the reaction volume (from 25 to 2 μl) does not negatively affect the quality of libraries obtained.

**Figure 3.**
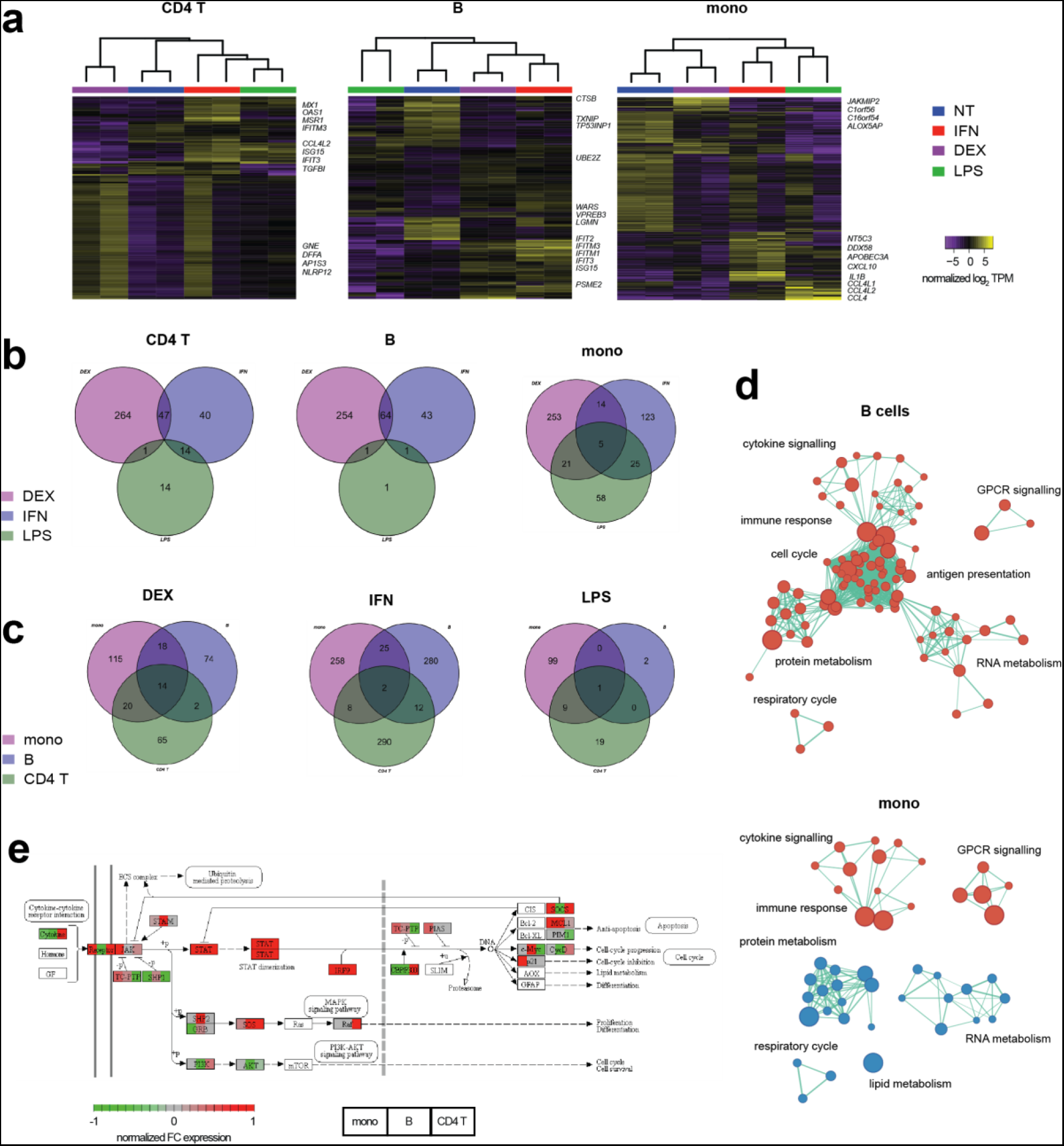
Genomic characterization of *ex vivo*-treated PBMCs. (**a**) Unsupervised clustering of untreated (NT) and treated (DEX, IFN, and LPS) PBMC subsets based on differentially expressed genes (FDR < 0.01). Top differentially expressed genes are labelled. Venn diagrams (**b, c**) showing common differentially expressed genes (FDR < 0.05) between treatments and subsets. (**d**) Gene set enrichment analysis (Reactome sets, FDR < 0.01) of IFN-treated B-cells and monocytes. Red nodes indicate upregulation, while blue nodes indicate down-regulation. Node sizes are proportional to the number of genes in the gene set, while edge lengths are inversely proportional to the number of overlapping genes between the sets. (**e**) Normalized fold-change in expression of Jak-STAT pathway genes in IFN-treated samples over untreated controls.

**Figure 4.**
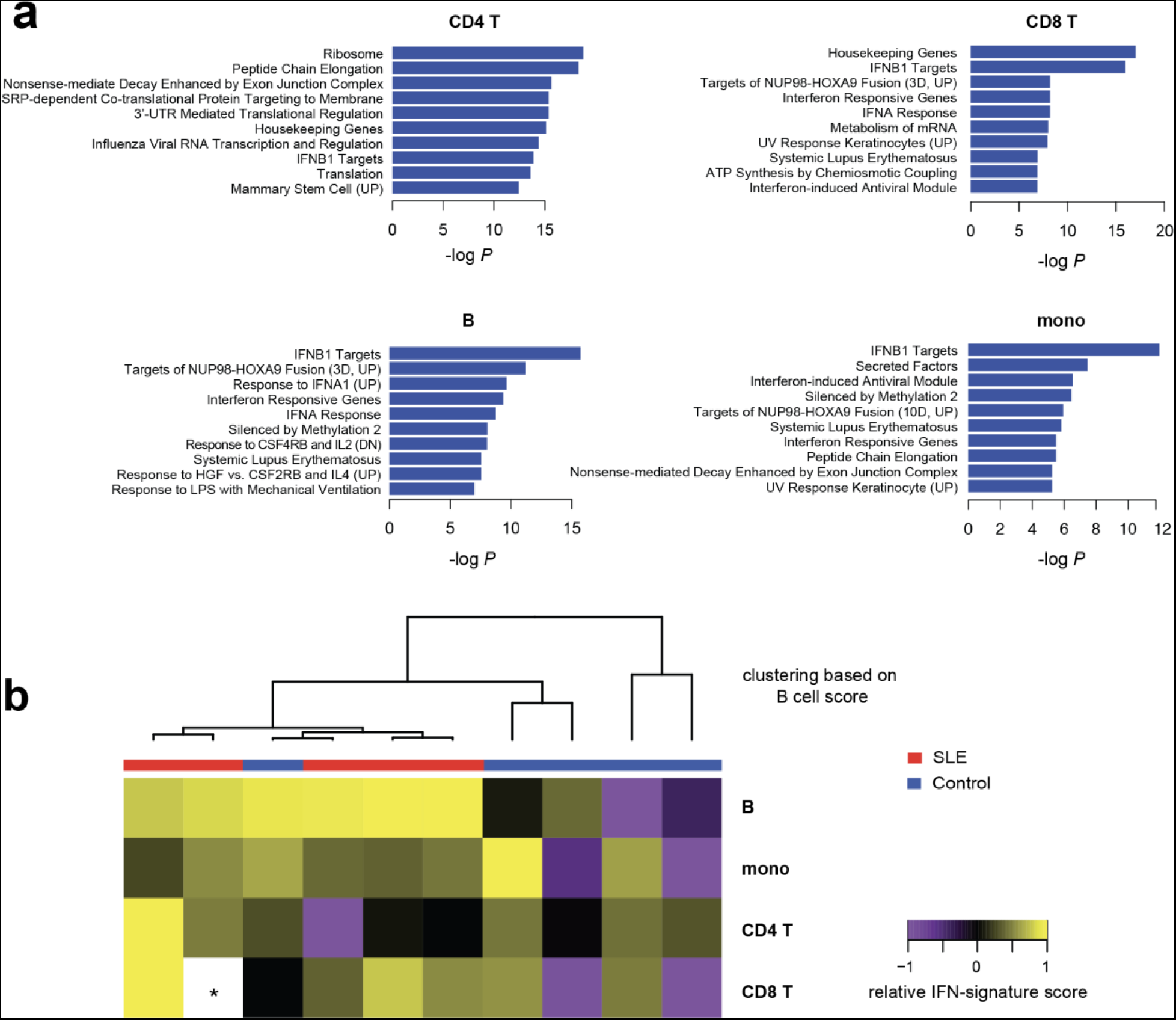
Transcriptional immune profiling of SLE patients. (**a**) Enriched gene sets (MSigDB C2) in SLE samples compared to healthy controls. *P-* values are adjusted for multiple gene set testing (Benjamini-Hochberg). (**b**) Heat map showing relative IFN-signature scores across different cell types of 10 patients. Scores (transcripts-per-million sum for 37 genes, **Supplementary Methods**) are mean-centered across each subset. Dendrogram shows clustering of patients based on IFN-signature scores for B cells. (*) Asterisk indicates missing data due to technical dropout.

**Table 1.**
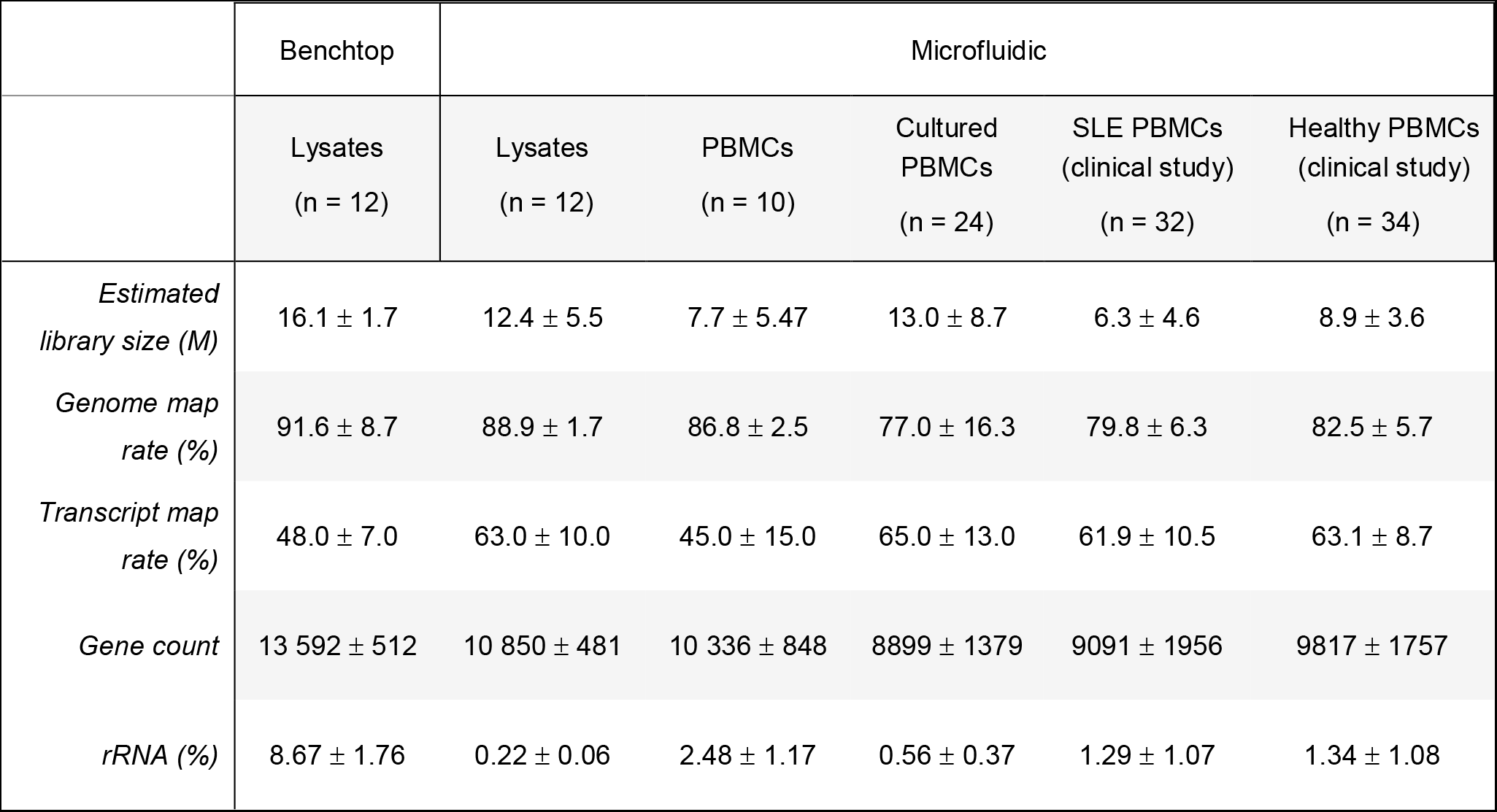
RNA-seq library statistics for the samples generated in this study. Two technical replicates are performed for each sample and isolated subset, *n* represents the total number of RNA-seq libraries generated for each column. Values are shown as mean ± S.D.

|  | Benchtop | Microfluidic |  |  |  |  |
| --- | --- | --- | --- | --- | --- | --- |
|  | Lysates<br>(n = 12) | Lysates<br>(n = 12) | PBMCs<br>(n = 10) | Cultured<br>PBMCs<br>(n = 24) | SLE PBMCs<br>(clinical study)<br>(n = 32) | Healthy PBMCs<br>(clinical study)<br>(n = 34) |
| *Estimated library size (M)* | 16.1 ± 1.7 | 12.4 ± 5.5 | 7.7 ± 5.47 | 13.0 ± 8.7 | 6.3 ± 4.6 | 8.9 ± 3.6 |
| *Genome map rate (%)* | 91.6 ± 8.7 | 88.9 ± 1.7 | 86.8 ± 2.5 | 77.0 ± 16.3 | 79.8 ± 6.3 | 82.5 ± 5.7 |
| *Transcript map rate (%)* | 48.0 ± 7.0 | 63.0 ± 10.0 | 45.0 ± 15.0 | 65.0 ± 13.0 | 61.9 ± 10.5 | 63.1 ± 8.7 |
| *Gene count* | 13 592 ± 512 | 10 850 ± 481 | 10 336 ± 848 | 8899 ± 1379 | 9091 ± 1956 | 9817 ± 1757 |
| *rRNA (%)* | 8.67 ± 1.76 | 0.22 ± 0.06 | 2.48 ± 1.17 | 0.56 ± 0.37 | 1.29 ± 1.07 | 1.34 ± 1.08 |

### Gene expression signatures of *ex vivo* stimulated PBMCs

We assessed whether our workflow can accurately and reproducibly profile the dynamic immune responses of different cell subsets. We cultured healthy PBMCs and applied three distinct treatments known to impact immune cells (LPS, IFN-α, and DEX) in duplicate. Using the microfluidic device to carry out multiplexed subset enrichments and RNA-seq library construction, we profiled the treatment response of three different subsets (CD4^+^T, B cells, and CD14^+^ monocytes). The device-processed libraries again showed strong reproducibility, co-clustering the duplicates based on differential gene expression responses (**Fig. 3a,Supplementary Fig. 7**) and accurately recording the differences in response between the three specific treatments used (**Fig. 3b, Supplementary Fig. 8**). Furthermore, our results highlight the heterogeneity in response between different cell subsets (**Fig. 3c-d**), as evidenced by the minimal overlap in differentially expressed genes and difference in enriched pathways between the profiled subsets. For B cells, IFN-α induces a proliferative response, as shown by the upregulation of cell cycle and metabolism pathways, while for monocytes, IFN-α induces an opposite effect. Even in the well-characterized Jak-STAT pathway, which is known to be directly activated by IFN-α, the pattern of downstream responses varied significantly across the three subtypes studied here (**Fig. 3e**). These results are consistent with previous reports that type I interferons can have either proliferative or suppressive effects on lymphocytes, depending on the relative timing of receptor co-activation^28^. In addition, these responses are greatly affected by cell-to-cell communication and the interplay between innate and adaptive immune activity^29–31^. This result emphasizes the importance of subset specific profiling to achieve higher resolution compared to monitoring immune responses compared with bulk expression profiling.

### Transcriptomic profiling of SLE patients

To demonstrate the utility of the device for disease studies, we profiled the immune state of 5 SLE patients and 5 healthy controls by isolating CD4^+^T, CD8^+^T, B cells, and CD14^+^ monocytes from cryopreserved PBMC samples (**Supplementary Table 1**). For each sample, 0.5 M cells were split into 8 channels on devices to isolate 4 subsets in duplicate and prepare RNA-seq libraries (**Supplementary Methods**). The data from our cohort validates the role of type I interferons in the pathogenesis of lupus^32^. We found type I interferon responses that are upregulated in SLE patients compared to matched healthy controls based on both differential expression and gene set analyses (**Figure 4a, Supplementary Fig. 9**). Interestingly, gene targets of the fusion protein NUP98-HOXA9, a potent driver of myeloid leukemia, were also enriched in all subsets. This supports previously published evidence that dysregulated lymphocyte proliferation is associated with both cancer and autoimmune disease, and could explain the increased malignancy risk in lupus patients^33^. Finally, to compare with previous gene expression studies, we generated an IFN-gene score based on a known panel of SLE signature genes previously identified in bulk studies^34^ (**Supplementary Methods, Supplementary Fig.10**). Our data shows that while this signature can be found across all the subsets we profiled, the difference between healthy and SLE scores is most pronounced in B cells (**Figure 4b**) (*P*= 0.05). This suggests that the diagnostic sensitivity and predictive power of the IFN signature for SLE may be improved by specifically profiling B cells instead of bulk PBMCs. Altogether, these initial findings show that gene expression responses in SLE differ across immune cell subsets and highlight the importance of subset-specific profiling in identifying disease signatures.

### DISCUSSION

Through our scalable microfluidic workflow, we demonstrate the utility of subset specific profiling of immune cells and its advantages over conventional bulk blood transcriptomics. Subset-specific analysis allows ready detection of biological signals from minority subsets by removing confounding effects from abundant cell populations such as the monocytes that dominate our test samples. Our method is complementary to the application of single cell transcriptomics approaches that are rising rapidly in popularity. For example, single-cell studies could point us to pathogenic subsets that can be enriched using the microfluidic device for large-scale research studies or clinical diagnostics. With this framework, scRNA-seq can be initially applied to a small cohort to identify clinically relevant subsets, after which, the integrated subset-specific microfluidic workflow can be used to scale-up to a larger cohort, increasing the study’s statistical power and lowering its cost. Another example would be the application of cell subset enrichment to help target cells of interest ahead of scRNA-seq. This type of workflow could dramatically improve the efficiency of scRNA-seq studies that target rare cell subsets by reducing the number of non-target cells that need be processed and sequenced to gain access to data from rare cells of interest.

In this report, we introduce the design and operation of a single microfluidic device that integrates both cell subset isolation and transcriptomic profiling. The device can reliably isolate cell subsets of interest and reproducibly construct RNA-seq libraries for next-generation sequencing. The sample multiplexing capability and free scaling of our microfluidic MACS implementation to low numbers of input cells while maintaining good enrichment performance are key advantages over conventional MACS approaches. Furthermore, the microfluidic device can be readily repurposed for other library preparation techniques, such as chromatin accessibility and DNA methylation profiling. We highlighted the importance of subset-specific profiling through *ex vivo* treatment of healthy PBMCs and the scalability of the workflow by profiling cell subsets in multiplex from SLE patient clinical samples. While the device does not yet reach the purity levels of FACS, the on-device enrichment approach boosts the signal from target subsets relative to bulk profiles by a significant 2- to 13-fold, enabling robust detection of weaker signals. This device will enable high-resolution monitoring of immune responses in clinical studies, especially in applications where blood samples or other inputs bear limited numbers of target cells, and large-scale immunomonitoring studies where significant sample throughput is required.

## METHODS

### Study samples

Human blood samples were obtained either from Research Blood Components (MA, USA) for technical validation experiments, or from collections at the Brigham and Women’s Hospital, MA USA (**Supplementary Table 1**). Research on the samples was approved by Institutional Review Boards at the Broad Institute of MIT and Harvard, MA (USA) and Brigham and Women’s Hospital, MA (USA). Blood samples from SLE patients and healthy control donors were drawn with EDTA Vacutainer tubes (BD Biosciences) and processed within 3 h of collection.

### Isolation of PBMCs from whole blood

Cells were isolated from whole blood samples using density gradient centrifugation. Whole blood was diluted 1:1 with 1X PBS, layered on top of Ficoll-Paque Plus (GE Healthcare), and centrifuged at 1200g for 20 min. The PBMC layer was retrieved, resuspended in 10 mL RPMI-1640 (Gibco), and centrifuged again at 300 g for 10 min. The cells were counted using a manual hemocytometer, resuspended in FBS (Gibco) with 10% DMSO (Sigma), and aliquoted in 1 mL cryopreservation tubes at a concentration of 5 M cells/mL. The tubes were kept at -80 °C overnight, then transferred to liquid nitrogen for long-term storage. Prior to processing, cells were thawed at 37 °C for 3 min, resuspended in 10 mL RPMI-1640 supplemented with 10% FBS (Gibco), and centrifuged at 300 g for 5 min. The cells were then resuspended in the desired concentration or buffer, depending on the experiment.

### Microfluidic device design and fabrication

The microfluidic device was fabricated using a previously published protocol^25^ with minor modifications. Flow layer molds were patterned in 3 steps: (1) rectangular 75 μm, (2) rectangular 200 μm, and (3) rounded 60 μm. All silicon wafers were pre-coated with hexamethyldisilazane (Sigma) before spin-coating. Rectangular features were prepared by spin-coating SU-8 2075 (Microchem) on a silicon wafer. The coated wafers were patterned by ultraviolet exposure (OAI 206 mask aligner) through a mask printed at 20,000 dpi (Fineline Imaging, design files are included in Supplementary Material). The features were then developed using SU-8 developer (Microchem). The rounded features were produced by spin coating AZ-40XT photoresist (Microchem), patterning the wafer with UV exposure and a mask, developing with AZ 400K developer (Microchem). After development, the wafer was subjected to an additional curing step (105° C for 10 min) to round the features. The control layer mold was patterned in one step: (1) rectangular 40 μm, using methods similar for the flow layer with SU-8 2015 photoresist (Microchem). Device production was carried out using standard soft-lithography, following the same published protocol, with the exception of final bonding to a silicon wafer substrate.

### Magnetic affinity cell isolation and microfluidic implementation

Magnetic affinity cell sorting was done using commercially available EasySep kits (CD14, CD19, CD4, CD8 positive isolation II and T cell negative isolation) from StemCell Technologies. In order to implement the isolation protocols on the device, the buffers were modified and volumes were scaled accordingly. EasySep buffer (StemCell Technologies) was supplemented with 10% FBS (Gibco) and 0.2% Pluronic-F127 (Sigma), in order to reduce non-specific cell adhesion in the PDMS channels. The micro-channels were also pre-incubated with 1% Pluronic-F127 (Sigma) prior to cell isolation. Neodynium magnets (Grainger) with a 43 lb pull were used for all magnetic capture steps.

### Flow cytometry and fluorescence-activated cell sorting

For assessment of isolation purities, flow cytometry was conducted using the Cytoflex system (Beckman Coulter). For RNA-seq library validation experiments and benchtop comparisons, PBMCs were sorted using the MoFlo Astrios (Beckman Coulter). Lysate pools were generated by sorting 5000 cells into 20 μl TCL buffer + 1 ul 20 mg/ml proteinase K (Qiagen) and stored at - 80 °C to maintain RNA integrity. The following panel was used for both purity assessment and sorting: DAPI, CD45 BV605, CD3 AF700, CD4 FITC, CD8 PE, CD14 APC, CD19 PE-Cy7, CD56 BV650 (all IgG1κ, BioLegend). Flow cytometry data was analyzed using FlowJo v10.1.

### Low-input RNA-seq, microfluidic implementation, and sequencing

RNA-seq was performed using Smart-Seq2^27^ with minor modifications. Cells were sorted into 19 μl TCL buffer + 1 ul 20 mg/ml proteinase K (Qiagen) and their RNA was purified by a 2.2x SPRI clean up with RNAClean XP magnetic beads (Agencourt) before reverse transcrition. For the microfluidic implementation of the protocol, Tween-20 (Teknova) was added to all reactions at a final concentration of 0.5%. For mRNA capture, a biotinylated oligo (/5BiosG/-AAGCAGTGGTATCAACGCAGAGTAC-30T-VN) (Integrated DNA Technologies) was attached to streptavidin magnetic beads (New England Biolabs) following the manufacturer’s protocol. The beads were then used to capture mRNA from the lysates, and were washed with 10 mM Tris-HCl pH 7.5, 0.15 M LiCl, 1 mM EDTA, 0.5% Tween-20. The beads were then resuspended in the reverse transcription mix, following the same steps as the published protocol. cDNA processed on the benchtop and microfluidic device were amplified for 18 and 22 cycles, respectively. After amplification and clean-up, libraries were quantified using a Qubit fluorometer (Invitrogen) and their size distributions were determined using the Agilent Bioanalyzer 2100. After normalizing the amplicon concentrations to 0.1-0.2 ng/mL, sequencing libraries were constructed using the Nextera XT DNA Library Prep Kit (Illumina), following the manufacturer’s protocol. All RNA-seq libraries were sequenced with 38x37 paired-end reads using a MiniSeq or NextSeq (Illumina).

### *Ex vivo* stimulation of PBMCs

Healthy PBMCs were resuspended in RPMI-1640 supplemented with 10% FBS and 1X penicillin-streptomycin (Gibco). Cells are cultured at a density of 1 M/mL and stimulated with LPS (5 μg/mL) (eBioscience), dexamethasone (100 nM) (Millipore), IFN- (250 U/mL) (Abcam), or no treatment. The cells were cultured for 24 hr at 37 °C in a 5% CO2 environment before processing through the microfluidic device.

### RNA-seq data analysis

RNA-seq libraries were sequenced to a depth of 5-15M reads per sample. All technical validation libraries were subsampled to 10 M reads to remove potential confounding effects of sequencing depth. Sequencing reads were aligned to the UCSC hg19 transcriptome using STAR^35^ and used as input to generate QC statistics with RNA-SeQC^36^. RSEM^37^ was used to generate an expression matrix for all samples. Both raw count and TPM (transcripts per million) data were analyzed using edgeR and custom R scripts. Lowly expressed genes with log2(CPM) less than 5 were filtered out before analysis. Gene set analyses were performed using the Kolmogorov–Smirnov test implementation in gage^38^. Cytoscape and the enrichMap^39^ module extension was utilized to visualize pathway-specific differential expression data.

## ACKNOWLEDGEMENTS

The authors thank Soohong Kim, Georgia Lagoudas, Anthony Kulesa, Navpreet Ranu, David Lieb, Arnon Arazi, and all the members of the Blainey and Hacohen Labs (Broad Institute) for helpful discussions. The authors also thank Samantha Leff for assistance in device fabrication, and the Broad Flow Cytometry core for assistance in the FACS experiments. This work was supported by grant U24 AI118668 from the National Institute of Allergy and Infectious Disease to N.H. P.C.B.). P.C.B was supported by a Career Award at the Scientific Interface from the Burroughs Welcome Fund. N.H. was supported by the David P. Ryan, MD Endowed Chair in Cancer Research.

## AUTHOR CONTRIBUTIONS

D.V. and P.C.B. designed the device. M.R., D.V., and K.B. fabricated microfluidic devices, implemented and optimized the microfluidic implementations of the MACS and RNA-seq protocols, and performed all device experiments. M.R. and T.E. performed experiments for benchtop protocol comparisons. M.R. analyzed all the RNA-seq data and designed the *ex vivo* experiments. T.E., P.H., and D.A.R. aided in clinical sample processing. N.H. and P.C.B. conceived and supervised the study. M.R. and P.C.B. prepared the manuscript, all authors reviewed and edited the final manuscript.

